# In-depth immunometabolic profiling by measuring cellular protein translation inhibition via bioorthogonal noncanonical amino acid tagging (CENCAT)

**DOI:** 10.1101/2023.08.03.551800

**Authors:** Frank Vrieling, Hendrik J.P. van der Zande, Britta Naus, Lisa Smeehuijzen, Bob J. Ignacio, Kimberly M. Bonger, Jan Van den Bossche, Sander Kersten, Rinke Stienstra

## Abstract

**Motivation:** Extracellular Flux (XF) analysis has been a key technique in immunometabolism research, measuring cellular oxygen consumption rate (OCR) and extracellular acidification rate (ECAR) to determine immune cell metabolic profiles. However, XF analysis has several limitations, including the need for purified adherent cells, relatively high cell numbers, and specialized equipment. Recently, a novel flow cytometry-based technique called SCENITH (Single Cell Energetic metabolism by profiling Translation inhibition) was introduced, which measures the inhibition of cellular protein synthesis as a proxy for metabolic activity in single cells. A limitation of this technique is its reliance on fluorescent staining of intracellular puromycin, a toxic antibiotic. To address this, we propose an alternative approach using biorthogonal noncanonical amino acid tagging (BONCAT) to measure protein synthesis.

**Summary:** The field of immunometabolism has revealed that cellular energy metabolism significantly contributes to immune cell function. Disturbances in immune cell metabolism have been associated with various diseases, including obesity, atherosclerosis, and cancer. To further advance immunometabolic research, developing novel methods to study the metabolism of immune cells in complex samples is essential. Here, we introduce CENCAT (Cellular Energetics through Non-Canonical Amino acid Tagging). This technique utilizes click-labeling of alkyne-bearing non-canonical amino acids (ncAAs) to measure protein synthesis inhibition as a proxy of metabolic activity. CENCAT successfully reproduced known metabolic signatures of immune cell activation. Specifically, LPS/IFNγ-induced classical activation increased glycolytic capacity, and IL-4-induced alternative activation enhanced mitochondrial dependence in human primary macrophages. The assay’s applicability was further explored in more complex samples, including peripheral blood mononuclear cells (PBMCs) from healthy volunteers, which revealed diverse metabolic rewiring in immune cell subsets upon stimulation with different activators. Finally, CENCAT was used to analyze the cellular metabolism of murine tissue-resident immune cells from various organs. Principal component analysis (PCA) revealed tissue-specific clustering based on metabolic profiles, likely driven by microenvironmental priming of tissue-resident immune cells. In conclusion, CENCAT offers valuable insights into immune cell metabolic responses and presents a powerful platform for studying immune cell metabolism in complex samples and tissue-resident immune populations in both human and murine studies.

## Introduction

The expanding field of immunometabolism has demonstrated that the functional properties of immune cells are dependent on their metabolic profile. When immune cells are activated, their energy demand increases to support proliferation and effector functions such as the production of cytokines and antibodies. This process requires a significant reprogramming of cellular metabolic pathways. Upon activation, many immune cell types switch from oxidative mitochondrial metabolism to glycolysis, a phenomenon known as the ‘Warburg effect’ as it was first described in tumor cells by Otto Warburg (1). Although essential for immune function, alterations in immune cell metabolism have been associated with various pathologies. For example, chronic low-grade inflammation in adipose tissue is linked to metabolic rewiring of adipose tissue macrophages in obesity (2), while ‘primed’ glycolytic inflammatory monocytes are associated with atherosclerosis (3). In tumors, immune cell metabolism is reprogrammed by limitations in nutrient availability and the accumulation of metabolic byproducts such as lactate, hindering their effector functions (4). Hence, technologies that enable detailed analysis of cellular metabolism are essential for studying the underlying mechanisms of inflammatory diseases.

Extracellular Flux (XF) analysis has been instrumental in making many discoveries in the immunometabolism field. This technique measures cellular oxygen consumption rate (OCR) and extracellular acidification rate (ECAR) to determine functional metabolic profiles. However, XF analysis has several limitations, including its reliance on purified, adherent cells, relatively high cell numbers, and specialized equipment and reagents. With the advent of single-cell technologies, developing novel methods to determine functional metabolic profiles of immune cells in complex samples is of great importance. A paper by Argüello et al. recently introduced a technique called SCENITH (Single Cell Energetic metabolism by profiling Translation inhibition). The core principle of SCENITH is that the rate of cellular protein synthesis is a surrogate for metabolic activity and can be used to determine cellular metabolic dependencies. In SCENITH, the incorporation of the antibiotic puromycin into nascent proteins is measured as a proxy for protein synthesis under conditions of metabolic inhibition (2-deoxy-D-glucose to inhibit glucose metabolism, oligomycin to inhibit mitochondrial ATP synthesis), allowing for the determination of cellular dependence on glucose and mitochondrial metabolism. Since SCENITH is a flow cytometry-based assay, it offers single-cell resolution and requires significantly fewer cells than XF analysis. This method has been compared and validated against Seahorse XF analysis (5, 6) and is thus a promising tool for determining functional metabolic profiles of immune cells in complex samples.

A significant limitation of using puromycin as a proxy for protein synthesis is its inherent toxicity, as it can induce ER stress (7) and other toxic effects (8). Additionally, puromycin was found to not accurately measure protein translation under energy-starved conditions such as 2-DG treatment, which could be problematic for its application in SCENITH (9). Therefore, we propose an alternative way of measuring protein synthesis in SCENITH through biorthogonal non-canonical amino acid tagging (BONCAT) (10). In BONCAT, cells are incubated with a non-canonical amino acid (ncAA) analog containing a minimal chemical modification, which can subsequently be targeted to fluorescently tag nascent proteins through copper(I)-catalyzed azide-alkyne cycloaddition (CuAAC) (11), more commonly known as the ‘click’ reaction. Among others, BONCAT has been used for affinity purification of nascent proteins (12), to track and localize the newly synthesized proteins in cells (13), and to measure SLC1A5-mediated amino acid uptake (14). A significant advantage of employing BONCAT for measuring protein synthesis is that ncAAs are non-toxic (15). The most widely used ncAAs are two methionine analogs: the azide-bearing azidohomoalanine (AHA) and the alkyne-bearing homopropargylglycine (HPG). As these require depletion of endogenous methionine for adequate protein labeling, a novel threonine ncAA analog, β-ethynylserine (βES), was recently developed to be able to tag proteins in complete growth medium (16).

Here, we investigated the potential of BONCAT using HPG and βES for measuring metabolic dependencies in multiple sample types, an approach we have termed CENCAT (Cellular Energetics through Non-Canonical Amino acid Tagging). We first show that cellular protein synthesis results in HPG accumulation over time. We subsequently validate our method compared to the original SCENITH protocol in a primary macrophage model of classical versus alternative activation. Next, we tested CENCAT using HPG or βES for metabolic profiling of peripheral blood mononuclear cells (PBMCs) under different stimulatory conditions. Finally, we demonstrate the technique’s applicability for the analysis of tissue-resident immune cell metabolism in mice.

## Materials & Methods

### Reagents, culture media, and supplements

L-homopropargylglycine (HPG), THPTA, AZDye 405 Azide Plus, and AZDye 488 Azide Plus were bought from Click Chemistry Tools (now Vector Laboratories). Lipopolysaccharide from Escherichia coli O55:B5 (LPS), oligomycin, 2-deoxy-D-glucose (2-DG), L-cystine dihydrochloride, RPMI 1640 medium (with sodium bicarbonate, without methionine, cystine and L-glutamine) and homoharringtonine were purchased from Merck. Fetal calf serum (FCS) was acquired from Biowest. RPMI 1640 medium (with sodium bicarbonate, without L-glutamine and HEPES), GlutaMAX™, and dialyzed FCS were bought from Thermo Fisher. Interferon γ (IFNγ) and interleukin-4 (IL-4) were acquired from Peprotech. ODN2006 was purchased from HyCult. Puromycin was obtained from InvivoGen. TransAct was bought from Miltenyi Biotec. Penicillin-Streptomycin Solution was obtained from Corning. β-ethynylserine-HCl (βES) was synthesized as described earlier (16) and dissolved in an equimolar NaOH solution to neutralize the pH. βES is available from Kimberly Bonger’s lab upon request.

### Flow cytometry antibodies and reagents

Anti-human antibodies: CD4-BV605, CD8-BV650, CD14-APC, CD16-PerCP-Cy5.5, CD19-PE-Dazzle594, CD45RA-BV785, CD62L-BV405 and HLA-DR-PE-Cy7 were bought from Biolegend. CD56-PE was purchased from Beckman Coulter. Anti-puromycin-Alexa Fluor 488 (clone R4743L-E8) was obtained by material transfer agreement (MTA) from Argüello et al. (6)

Anti-mouse antibodies: B220-BV510, CD4-FITC, CD8a-PE-Cy7, CD11b-BV650, CD11c-BV605, CD45-PerCP-Cy5.5, CD64-APC, F4/80-FITC, Ly6C-Alexa Fluor 700, MHCII-BV785, Siglec-F-PE-Dazzle594 and TIM4-PE were acquired from Biolegend. CLEC2-FITC was purchased from Bio-Rad.

Zombie Aqua and Zombie NIR Fixable Viability Dyes, as well as Human and Mouse TruStain FcX Fc Receptor Blocking Solution, were from BioLegend. CF700-SE and CF750-SE were bought from Biotium. Brilliant Stain Buffer Plus (BD Biosciences) was added to antibody mixes to prevent staining artifacts caused by interactions between Brilliant Violet dyes. In mouse experiments, True-Stain Monocyte Blocker (Biolegend) was used to avoid non-specific antibody binding of PE- and APC-based tandem dyes by myeloid cells.

### Peripheral blood mononuclear cell (PBMC) isolation and stimulation

EDTA blood was collected from healthy volunteers after acquiring informed consent as per the norms of the International Declaration of Helsinki. Blood tubes were pooled and diluted 1:1 in PBS, after which PBMCs were isolated by Ficoll Paque Plus (Merck) gradient centrifugation in Leucosep™ tubes (Greiner Bio-One). PBMC layers were collected in a 50 ml tube and washed thrice with PBS before counting using a hemocytometer. PBMCs were resuspended at a density of 10×10^6^ cells/mL in RPMI 1640 medium supplemented with 10% FCS, GlutaMAX, and P/S and cultured in FACS tubes at 37°C/5%CO_2_ while shaking (100 RPM). To activate PBMCs, cells were stimulated with LPS (100 ng/mL), ODN 2006 (1 μM), TransAct (1:100), or a mixture of all three stimuli for 2 hours in total.

### Human monocyte isolation and macrophage differentiation

Human primary monocytes were isolated from buffy coats of healthy blood bank donors through magnetic-activated cell sorting (MACS). After PBMC isolation as described above, CD14^+^ monocytes were magnetically labeled using MojoSort Human CD14 Nanobeads (Biolegend) and subsequently separated on LS columns using a QuadroMACS™ Separator (Miltenyi Biotec). Isolated monocytes were counted using a Vi-CELL XR Cell Analyzer (Beckman Coulter), resuspended at a density of 1×10^6^ cells/mL in RPMI 1640 medium supplemented with 10% FCS, GlutaMAX and P/S, and seeded in T75 flasks using 10 mL per flask (10×10^6^ cells). Monocytes were cultured for six days at 37°C/5%CO_2_ in the presence of either M-CSF (50 ng/mL; Miltenyi Biotec) or GM-CSF (5 ng/mL; Miltenyi Biotec) for macrophage differentiation. At day 3 of differentiation, 5 mL of fresh medium containing M-CSF or GM-CSF was added to each flask.

After differentiation, macrophages were harvested by trypsinization and seeded in 24-well plates at a density of 3×10^5^ cells/well (GM-CSF) or 4×10^5^ cells/well (M-CSF). Macrophages were subsequently left untreated, classically activated by a combination of LPS (100 ng/mL) and IFNγ (10 ng/mL), or alternatively activated using IL-4 (20 ng/mL) for 24 hours.

### Mice experiments

All experiments followed the Guide for the Care and Use of Laboratory Animals of the Institute for Laboratory Animal Research and were approved by the Central Authority for Scientific Procedures on Animals (CCD, AVD10400202115283) and the Institutional Animal Care and Use Committee of Wageningen University. Tissues were collected from mice as part of experimental protocols 2021.W-0016.008 and 2021.W-0016.007. Naïve, 12-16-week-old, male wild-type C57BL/6J mice were sacrificed by cervical dislocation, and epidydimal white adipose tissue (eWAT) fat pads, kidneys, liver, lungs, and spleen were collected in RPMI for tissue-resident immune cell isolation. Before collecting tissues, the peritoneal cavity was washed with 10 mL ice-cold PBS supplemented with two mM EDTA to isolate peritoneal exudate cells (PECs). Tissue-resident immune cells were isolated as described previously (17, 18) with minor adaptations. Briefly, eWAT, kidneys, liver, and lungs were cut into small pieces using razors, and the spleen was mechanically disrupted using a syringe plunger. eWAT fat pads were digested in 5 mL digestion buffer containing 1 mg/mL collagenase type II from Clostridium histolyticum (Sigma-Aldrich) in Krebs buffer supplemented with 100 mM HEPES, 20 mg/mL BSA and 6 mM D-Glucose for 45 minutes at 37°C/5%CO_2_ while shaking (100 RPM). Digestion was stopped by adding 5 mL wash buffer (PBS supplemented with 1% FCS and 2.5 mM EDTA), and the solution was filtered through 250 μm Nitex filters (Sefar). Infranatant containing SVF was collected, and erythrocytes were lysed using ice-cold erythrocyte lysis buffer (in-house, 0.15 M NH_4_Cl, 1 mM KHCO_3_, 0.1 mM Na_2_EDTA). Cells were finally filtered through 40 μm cell strainers (PluriSelect).

Kidneys and lungs were digested in 5 mL digestion buffer containing 1 mg/mL collagenase V, 1 mg/mL dispase II, and 30 U/mL DNase I in RPMI for 30 minutes at 37°C/5%CO_2_ while shaking (100 RPM). Digestion was stopped by adding 5 mL wash buffer, and the digest was filtered through 100 μm cell strainers (PluriSelect). After erythrocyte lysis and filtering through 40 μm cell strainers, leukocytes were purified using magnetic-assisted cell sorting (MACS) using CD45 microbeads (35 μl per sample, Miltenyi Biotec) according to manufacturer’s instructions.

The liver was digested in 5 mL RPMI supplemented with 1 mg/mL collagenase V, 1 mg/mL dispase II, 1 mg/mL collagenase D, and 30 U/mL DNase I for 25 minutes at 37°C/5%CO_2_ while shaking (100 RPM). Digest was filtered through 100 μm cell strainers and washed twice with 40 mL wash buffer (300 RCF, 5 minutes, 4°C). After erythrocyte lysis and filtering through 40 μm cell strainers, leukocytes were isolated using CD45 MACS, similar to kidneys and lungs.

The spleen was digested in 5 mL digestion buffer containing 1 mg/mL collagenase D and 30 U/mL DNase I in RPMI for 30 minutes at 37°C/5%CO_2_ while shaking (100 RPM). Digest was filtered through 100 μm filters, incubated with erythrocyte lysis buffer, and filtered again through 40 μm filters.

All isolated and filtered cells were counted using a hemocytometer. eWAT, kidney, and peritoneal leukocytes were split over four wells, 2.5-5x10^5^ liver and lung leukocytes/well, and 1x10^6^ splenocytes/well were plated in 96-well U-bottom plates for CENCAT.

### CENCAT

#### Methionine depletion for experiments with HPG

Cells (human PBMCs, human primary macrophages) incubated with HPG were first cultured for 30-45 minutes at 37°C/5%CO_2_ in methionine-free RPMI 1640 supplemented with 65 mg/L L-cystine dihydrochloride, 10% dialyzed FCS, GlutaMAX and P/S to deplete intracellular methionine levels. Any ongoing stimulation (e.g. LPS, ODN 2006, TransAct) was refreshed in this medium.

#### Metabolic inhibitor and ncAA incubation

Cells were pre-incubated with metabolic inhibitors (10× work stocks in complete RPMI) for 15 minutes to ensure complete shutdown of glucose and/or mitochondrial metabolism. Methionine-free RPMI was used for HPG experiments. The following four treatments were applied in technical duplicate (human PBMCs, human primary macrophages) or uniplo (mouse tissue-resident immune cells) per experimental condition: DMSO (control), 2-deoxyglucose (2-DG; 100 mM), oligomycin (1 μM), and a combination of 2-DG and oligomycin. All treatments were corrected for DMSO and H_2_O content. Cells were subsequently treated with HPG (100 μM) or βES (500 μM) and incubated for 30 minutes at 37°C/5%CO_2_. After ncAA incorporation, macrophage samples were washed 1× with PBS, harvested by scraping, and transferred to a V-bottom 96-well plate. PBMC samples and mouse tissue-resident immune cells were transferred to a V-bottom 96-well plate and pelleted through centrifugation at 500RCF for 3 minutes, after which the supernatant was discarded by firmly flicking the plate once. Cells were then washed 1× with PBS. For samples stained according to the original SCENITH protocol, puromycin (end concentration 10 μg/mL) was added instead of ncAAs.

HPG was added directly after methionine depletion with or without homoharringtonine (20 μg/mL) or DMSO control for kinetic experiments. Macrophages were harvested at T = 0, 0.5, 1, 2, 4, and 6 hours post-treatment.

#### Fluorescent labeling of nascent proteins and flow cytometry

Human cells were pelleted by centrifugation and incubated with Zombie Aqua Fixable Viability Dye (1:1000 in PBS) for 5-10 minutes at RT in the dark. Cells were then pelleted by centrifugation and fixed in 2% formaldehyde for 15 minutes at RT. After fixation, cells were pelleted and washed 1× with PBS before permeabilization with 0.1% saponin in PBS/1%BSA or 1X permeabilization buffer (Invitrogen) for 15 minutes at RT. Next, cells were washed 1× with PBS and barcoded with amine-binding dyes (CF700- and CF750-succinimidyl esters, 1 mM stocks) for 5 minutes at RT. Barcoding dyes were applied in four combinations of two dilutions (1:1000 and 1:100,000), allowing for pooling the four treatment conditions per replicate. Cells were then washed 1× with PBS, and barcoded samples were pooled in Click buffer (100 mM Tris-HCl, pH 7.4). ncAAs were labeled through copper(I)-catalyzed azide-alkyne cycloaddition (CuAAC) using the following reaction mix in Click Buffer: 0.5 mM Cu(II)SO_4_, 10 mM sodium ascorbate, 2 mM THPTA and 0.5 μM AF488 Azide Plus. Freshly prepared sodium ascorbate solution (100 mM stock) and AZDye 488 Azide Plus were added to the mixture just before addition to the samples. Click reaction was performed for 30 minutes at RT, after which the cells were washed 2× with FACS buffer (PBS/1%BSA + 2 mM EDTA). For samples stained according to the original SCENITH protocol, Alexa Fluor 488-conjugated anti-puromycin antibody diluted 1:100 in FACS buffer was added instead of the click reaction mix. PBMC samples were then blocked with Human TruStain FcX Fc Receptor Blocking Solution (1:100) before staining with fluorescent antibodies for 15 minutes at RT in the presence of Brilliant Stain Buffer.

Mouse cells were processed similarly with minor adjustments. Mouse cells were not barcoded, 0.5 μM AF405 Azide Plus was used for CuAAC, and mouse tissue-resident immune cells were stained with a fluorescent antibody cocktail in the presence of Brilliant Stain Buffer and TrueStain Monocyte Blocker.

Cells were washed 1× with FACS buffer after staining and acquired on a CytoFLEX S cytometer (Beckman Coulter).

#### Flow cytometry and statistical analysis

SCENITH parameters were calculated as described previously by Argüello et al. (6).

Human macrophage and murine tissue flow cytometry data were analyzed using FlowJo software version 10.8.1 (Becton Dickinson). Human PBMC flow cytometry data were analyzed using OMIQ software from Dotmatics (www.omiq.ai, www.dotmatics.com).

Principal component analysis (PCA) was performed in R version 4.2.2 using the mixOmics package version 6.23.4 (19). R plots were made using the following packages: ggplot2 version 3.4.22 (20), cowplot version 1.1.1 (21), and ggh4x 0.2.4 (22). Statistical significance was tested as indicated by Repeated One-Way ANOVA with Dunnett’s multiple comparisons test, Two-Way ANOVA with Sidak correction for multiple testing or paired t-test using GraphPad Prism software version 8.01.

## Results

### CENCAT: assay concept

The assay concept is depicted in Figure 1. After sample workup and plating (1), cells are pre-incubated with DMSO as control, 2-DG to inhibit glucose metabolism, oligomycin to inhibit mitochondrial respiration, or the compounds in combination to block ATP production completely (2). An alkyne-bearing ncAA is subsequently added as a substrate for protein synthesis (3). Cells are then fixed and permeabilized, after which intracellular ncAA-tagged proteins can be labeled through click reaction with a fluorescent azide probe to finally be measured through flow cytometry (4). The relative cellular metabolic dependence on glucose and mitochondrial respiration is calculated by comparing ncAA mean fluorescence intensities between the different inhibitor conditions (5). Glucose dependence is calculated by dividing the mean fluorescence intensity difference (ΔMFI) between DMSO and 2-DG by the ΔMFI between DMSO and 2-DG + oligomycin. The inverse of glucose dependence gives you the fatty acid/amino acid oxidation (FAO/AAO) capacity. Mitochondrial dependence is the ΔMFI between DMSO and oligomycin divided by the ΔMFI between DMSO and 2-DG + oligomycin. The inverse of mitochondrial dependence is the glycolytic capacity.

**Figure 1:**
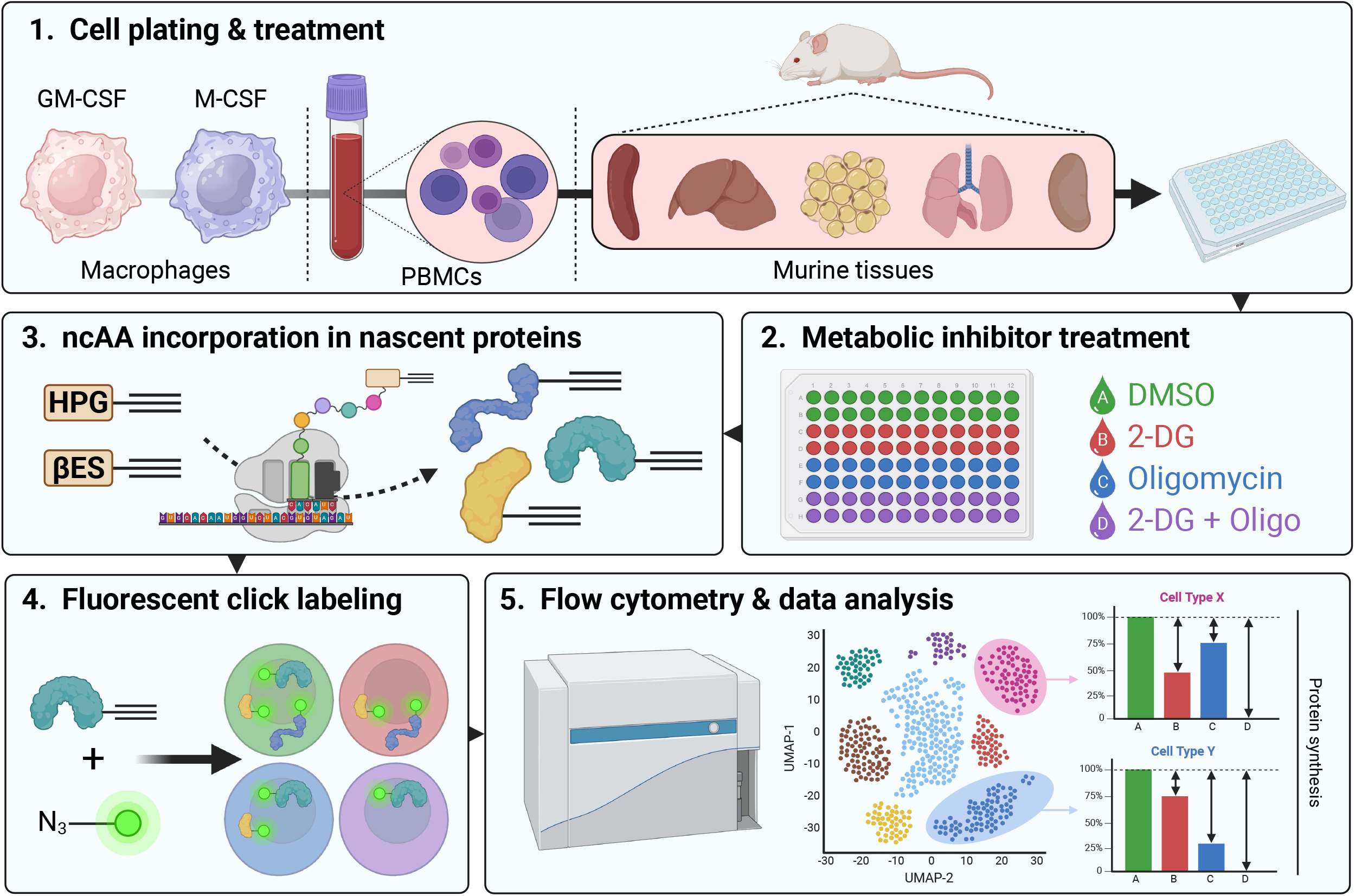
SCENITH-BONCAT assay concept. Created using BioRender (www.BioRender.com).

### HPG incorporation is a proxy of protein synthesis and can be used as an alternative to puromycin

HPG is an alkyne-bearing methionine analog (Figure 1A) previously used to tag nascent proteins (12). To validate whether HPG incorporation in immune cells reflects protein synthesis, M-CSF differentiated primary human macrophages were first methionine-depleted for 1 hour and subsequently treated with HPG for up to 6 hours in the presence or absence of homoharringtonine, a protein translation inhibitor. HPG accumulated in macrophages over time as measured by flow cytometry (Figure 2B). This accumulation was entirely blocked by treatment with homoharringtonine (Figure 2C), demonstrating that HPG incorporation can be used as a proxy for protein synthesis.

**Figure 2:**
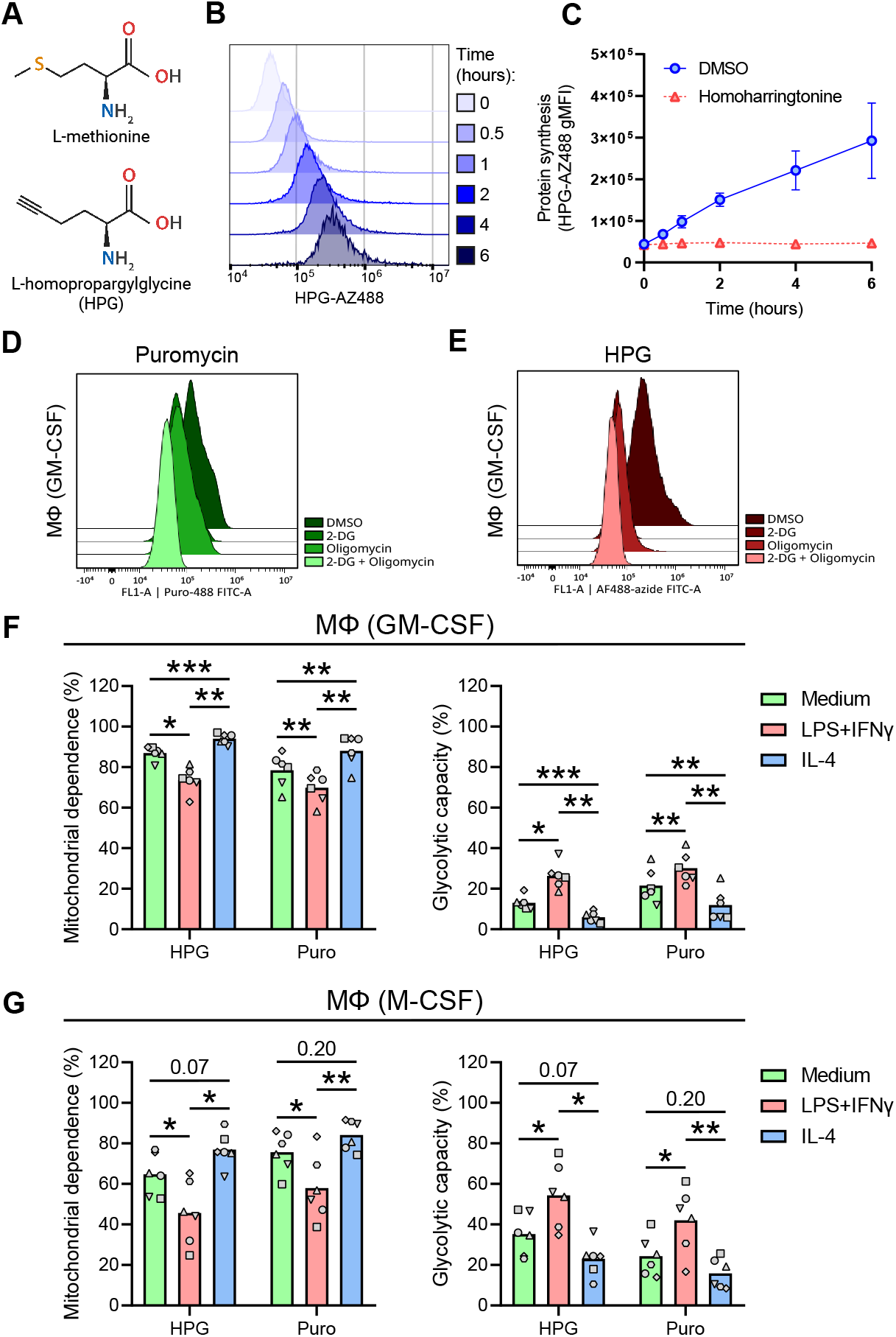
Comparison of CENCAT with the original SCENITH protocol. (A) Structures of L-methionine and HPG. (B-C) Primary human macrophages (M-CSF) were treated with HPG and harvested at different time points for click labeling. (B) Representative flow cytometry histograms of HPG-AZ488 staining at 0, 0.5, 1, 2, 4, and 6 hours post-treatment. (C) HPG incorporation kinetics of macrophages treated with homoharringtonine (red) or DMSO control (blue). Data are displayed as means ± SD of the geometric MFI (gMFI, n=3). (D-E) Representative histograms of protein synthesis for DMSO, 2-DG, Oligomycin, and 2-DG + Oligomycin conditions as measured using puromycin (D) or HPG (E) in GM-CSF macrophages. (F-G) Primary human macrophages (GM-CSF and M-CSF) were stimulated for 24 hours with culture medium (green), LPS + IFNγ (red), or IL-4 (blue) before SCENITH analysis using either HPG or puromycin (Puro) as substrates. (F) Mitochondrial dependence (%) and glycolytic capacity (%) of GM-CSF macrophages. (G) Mitochondrial dependence (%) and glycolytic capacity of M-CSF macrophages. Data are displayed as mean percentages ± SD (n=6). Significance was tested by Two-Way ANOVA with Sidak correction for multiple testing. Individual donors are displayed by different symbols. * = *p* < 0.05, ** = *p* < 0.01, *** = *p* < 0.001

Next, we tested whether CENCAT could reproduce the results of the original SCENITH protocol using the anti-puromycin antibody clone R4743L-E8. Both assay variants were applied side-by-side to analyze the metabolic profiles of primary human macrophages after classical activation (LPS + IFNγ) versus alternative activation (IL-4). Representative histograms of protein synthesis for the different inhibitor conditions are shown in Figure 1D for puromycin and Figure 1E for HPG. LPS activation in macrophages is associated with the upregulation of glycolysis and a disrupted tricarboxylic acid (TCA) cycle, whereas IL-4 activation is supported by enhanced mitochondrial oxidative phosphorylation (OXPHOS) (23). As expected, classical activation resulted in a relative increase in glycolytic capacity over mitochondrial dependence. At the same time, the inverse was observed for alternative activation in both GM-CSF (Figure 2F) and M-CSF (Figure 2G) differentiated macrophages. This result was obtained irrespective of whether HPG or puromycin detection was used as a proxy for protein synthesis. Both GM-CSF and M-CSF macrophages were highly glucose-dependent (>73%) across the different polarized states, as evidenced by both techniques (Figure S1A-B). The absolute values for glucose dependence were not significantly different between both techniques (Figure S1A). These results establish CENCAT as a viable method for performing immune cell metabolic profiling.

### CENCAT reveals differential metabolic responses to inflammatory stimulation in PBMCs

An essential improvement of SCENITH over other techniques, such as XF analysis, is its ability to perform metabolic profiling of multiple cell types simultaneously in complex samples. Therefore, we next tested the applicability of HPG-based CENCAT to determine metabolic dependencies in PBMCs using a nine-marker flow cytometry panel to identify B cells, monocytes, CD4 T cells, CD8 T cells, and NK cells (Figure 3B, and Figure S2 for gating strategy). We aimed to investigate the metabolic profile of these cells in both their resting state and after immune cell activation. A combination of immunological stimuli was applied to achieve this, as different immune cell types are activated through distinct cell surface receptors. For instance, myeloid cells, such as monocytes, highly express the TLR4 receptor, which recognizes LPS. B cells respond strongly to DNA containing unmethylated CpG sequences via TLR9 (24, 25), whereas T cells are activated by co-activating the T cell co-receptor CD3 and its co-stimulatory receptor CD28. Therefore, to determine the effect of immune activation on metabolic dependencies, PBMCs were either left untreated (Medium) or stimulated for 2 hours with LPS (TLR4 ligand), ODN2006 (CpG, TLR9 ligand), TransAct (synthetic CD3/CD28 agonist) or a mixture containing all three ligands (Figure 3A).

**Figure 3:**
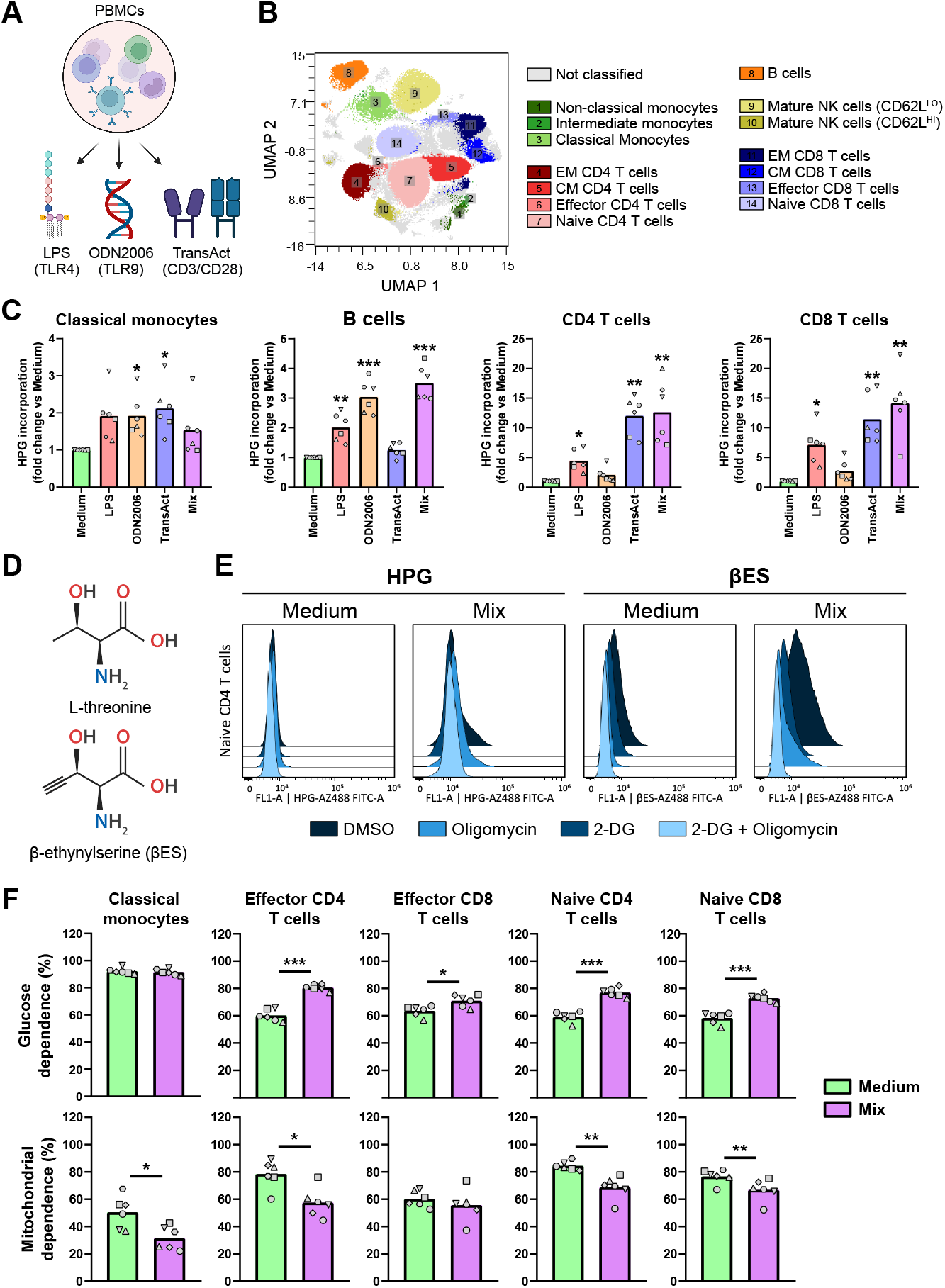
CENCAT analysis of PBMC metabolic profiles after different stimulations. (A) PBMCs isolated from healthy blood donors (n=6) were stimulated for 2 hours with Medium control (green), LPS (red), ODN2006 (orange), TransAct (blue), or all stimuli combined (Mix, purple). CENCAT was performed using HPG (C, E) G or βES (E, F). (B) Representative UMAP plot of PBMCs stained with 9-marker flow cytometry panel. Colors and numbers indicate different cell populations. (C) Fold changes of HPG incorporation compared to Medium control of classical monocytes, B cells, CD4 T cells, and CD8 T cells. Significance was tested by Repeated One-Way ANOVA with Dunnett’s multiple comparisons test. Individual donors are displayed by different symbols. * = *p* < 0.05, ** = *p* < 0.01, *** = *p* < 0.001. (D) Structures of L-threonine and βES. (E) Representative histograms of protein synthesis for DMSO, 2-DG, Oligomycin, and 2-DG + Oligomycin conditions as measured using HPG or βES in Naïve CD4 T cells. (F) Glucose dependence (%) and mitochondrial dependence (%) of classical monocytes, effector CD4 T cells, effector CD8 T cells, naïve CD4 T cells, and naïve CD8 T cells. Significance was tested by paired t-test. Bars indicate mean percentages and individual donors are displayed by different symbols. * = p < 0.05, ** = p < 0.01, *** = p < 0.001

HPG incorporation was differentially elevated by stimulation in all tested immune cells (Figure 3C). The stimuli mixture (Mix) generally yielded similar results as the most potent single stimulation for each cell type. As expected, B cells showed the highest HPG incorporation after treatment with the TLR9 ligand ODN2006, while both CD4 and CD8 T cells most strongly responded to CD3/CD28 activation by TransAct. CD14^+^/CD16^-^ classical monocytes showed a similar increase in HPG levels for all single stimulations, most likely as a result of paracrine signaling by other PBMCs. Noticeably, our results showed that HPG incorporation levels were low in several cell types. This was most apparent in untreated T cells, indicating that these cell types are relatively quiescent when inactive. (Figure S3A). As a result, small fluctuations in HPG signal skew the relative metabolic dependencies in these cells, leading to high variability (Figure S3B-C). Additionally, oligomycin treatment occasionally increased protein translation in T cells, leading to negative values for mitochondrial dependence. As HPG incorporation is known to be inefficient relative to methionine (26), we substituted HPG with βES (Figure 3D), a novel threonine-derived ncAA which reportedly has a ∼12.5-fold higher incorporation rate compared to HPG (16).

The use of βES as a substrate for CENCAT was analyzed in PBMCs left either unstimulated (Medium) or treated with the three ligand stimulation mixture (Mix) for 2 hours. βES displayed improved incorporation in T cells compared to HPG, as is illustrated by the flow cytometry histograms of Naïve CD4 T cells in Figure 3E. Only a low HPG signal was detected in the control condition (ΔMFI DMSO vs 2-DG + Oligomycin: 937 ± 578), which was elevated after stimulation (ΔMFI: 6547 ± 4139). Incorporation was significantly higher in naïve CD4 T cells incubated with βES (Medium control ΔMFI: 6715 ± 3909, p = 0.035 vs HPG; Stimulation Mix ΔMFI: 12865 ± 6629, p = 0.018 vs HPG). Furthermore, unlike HPG, βES-treated cells did not display any negative metabolic dependencies (Figure 3F, Figure S4C-D).

PCA analysis showed a clear group separation between samples stimulated with the stimulation mixture or medium control (Figure S4A). The top loadings of the first component (PC1) show that metabolic dependencies of classical monocytes and CD45RA^+^ T cells, comprising the effector and naïve subsets, were most important for group separation along this axis (Figure S4B). Stimulation significantly decreased mitochondrial dependence in these cell types except for effector CD8 T cells (Figure 3F), indicating a metabolic switch to glycolysis reminiscent of the Warburg effect. In line with this, activated CD45RA^+^ T cell subsets displayed a significantly increased dependence on glucose, whereas glucose dependence was already high (>90%) in classical monocytes at basal conditions. This increased reliance on glucose was also observed in most CD45RA^-^ central/effector memory T cell subsets (Figure S4C); however, these cells did not display a relative glycolytic shift in response to stimulation. Compared to classical monocytes, the non-classical and intermediate subsets showed a similar high dependence on glucose at baseline (Figure S4D) but a much higher mitochondrial dependence (90-98%). Unfortunately, these subsets could not be detected after stimulation. B cells and mature NK cells did not show a shift in metabolism in response to the stimulation mixture.

Altogether, owing to increased incorporation and, as a result, improved resolution of protein synthesis, we propose the use of βES over HPG for metabolic profiling of complex samples containing relatively quiescent cells such as naïve T lymphocytes.

### Metabolic profiling of murine tissue-resident immune cell populations

We next tested the potential of CENCAT to determine metabolic profiles of tissue-resident immune cell populations in mice. Immune cells from epididymal white adipose tissue (eWAT), kidneys, liver, lungs, peritoneal exudate cells (PEC) and spleen were isolated from wild-type C57BL/6J mice and subjected to CENCAT with βES. A 12-marker flow cytometry was used to identify conventional dendritic cell subsets (cDC1 and cDC2), tissue-specific macrophage populations, monocytes, B cells, neutrophils, eosinophils, CD4, and CD8 T cells. The complete gating strategies are depicted in Figure S5.

βES incorporation levels were variable across tissues and cell types (Figure 4A). In general, tissue-resident immune cells in PEC and spleen samples showed the highest βES fluorescence intensities, while βES accumulation was relatively low in kidney immune cells. Within tissues, DCs and macrophages often displayed the highest βES signal. βES incorporation and metabolic profiles could not be assessed in eosinophils, as these cells acquired a very high non-specific background staining after the fluorescent click reaction, even in the absence of βES (data not shown).

**Figure 4:**
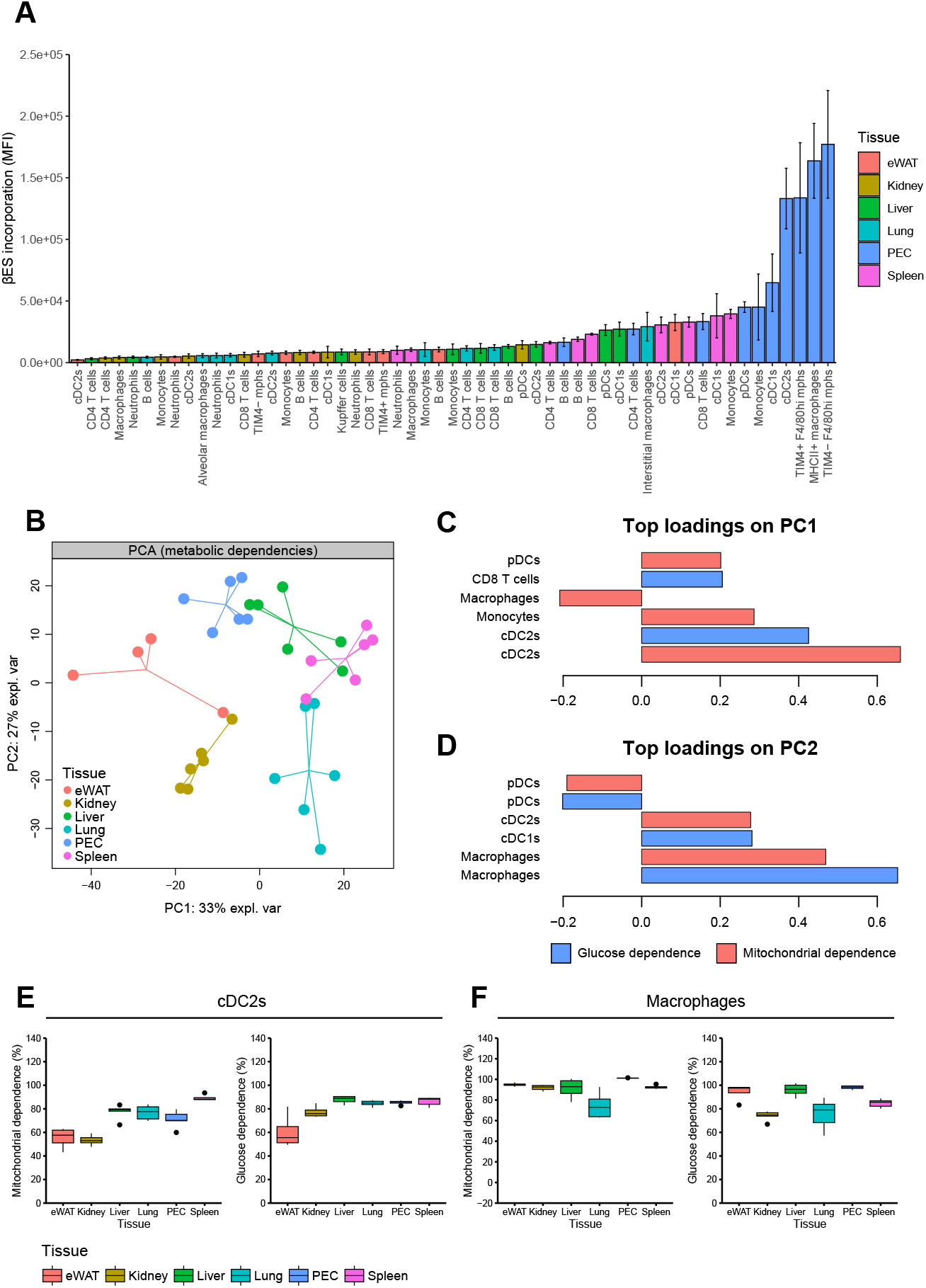
CENCAT analysis of murine tissue-resident immune cell populations. The following tissues were isolated from male C57BL/6J mice and subjected to CENCAT analysis: eWAT (red, n=4), kidney (yellow, n=6), liver (green, n=6), lung (cyan, n=6), PEC (blue, n=6), spleen (pink, n=6). (A) βES incorporation (MFI) of immune cell populations from all six tissues. (B) PCA score plot based on metabolic dependencies of immune cell populations in all six tissues. (C-D) Top loadings on PC1 (C) and PC2 (D) of the PCA score plot. Measures of glucose dependence are represented by blue bars and mitochondrial dependence by red bars. (E-F) Mitochondrial and glucose dependence (%) of cDC2s (E) and macrophages (F) from all six tissues.

To visualize potential differences in immune cell metabolic profiles across tissues, glucose and mitochondrial dependencies of shared immune populations were analyzed by PCA. Samples derived from the same tissue clustered together, and clear group separations were observable between different tissues (Figure 4B). The top loadings of the first and second components (PC1 and PC2), which respectively explain 33% and 27% of data variance, show that metabolic dependencies of myeloid cells were most important for projection along the axes, in particular cDC2s (Figure 4C) and macrophages (Figure 4D), indicating that these cells are most metabolically variable across different tissues. cDC2s from eWAT and kidney showed low levels of mitochondrial and glucose dependence compared to the other tissues, while spleen cDC2s were highly reliant on glucose and mitochondria (Figure 4E).

Further analysis of macrophage subsets showed that interstitial macrophages, Kupffer cells, and adipose tissue macrophages were similarly highly dependent on glucose and oxidative metabolism (Figure S6A-B). Within PEC, a subdivision in mitochondrial and glucose metabolism could be observed between MHCII+ monocyte-derived immature macrophages and F4/80^HI^ mature tissue-resident macrophages, which is in line with a recent study (27), which employed CENCAT with HPG. Neutrophils were the most glycolytic of all investigated cell types, particularly in adipose tissue, whereas monocytes, pDCs, cDC1s, B cells, and T cells primarily depended on oxidative metabolism (Figure S6C). Additionally, neutrophils and monocytes were highly reliant on glucose across all tissues, while this was more variable for pDCs, B cells, and T cells (Figure S6D).

## Discussion

Here, we established CENCAT as a valid approach for conducting in-depth metabolic profiling. Our findings demonstrate that ncAA incorporation can serve as an alternative to puromycin immunostaining as a proxy for protein synthesis. Among the tested ncAAs, the novel threonine analog βES emerges as the preferred choice over HPG, owing to its relatively high incorporation efficiency in metabolically inactive cells. We have shown that CENCAT is applicable for studying immune cells in peripheral blood mononuclear cells (PBMCs) and tissue-resident immune cells from murine tissues. This underscores its significance as a potent metabolic profiling tool in the immunologist’s arsenal.

The use of ncAAs over puromycin as a substrate to measure protein synthesis has several advantages. Firstly, puromycin is a toxic antibiotic often used as a selection agent for eukaryotic cells. Puromycin inhibits protein translation, leading to the formation of prematurely terminated truncated polypeptides (28). This apparent toxicity limits the maximal substrate incorporation time and could lead to unwanted effects. ncAAs, on the other hand, are incorporated into full-length proteins and are therefore reported to be non-toxic (15). Secondly, puromycin labeling was found to be inappropriate for accurately measuring overall protein synthesis rates during energy-starved conditions compared to the methionine-analog AHA, particularly in response to glucose starvation and 2-DG treatment (9). This could explain differences in glucose dependence measurements between both technologies.

Our CENCAT approach using HPG, similar to the original SCENITH assay using puromycin, showed that LPS/IFNγ-primed classical activation of monocyte-derived macrophages resulted in a relative increase in glycolytic capacity, while IL-4-induced alternatively activated macrophages displayed enhanced mitochondrial dependence. These results align with previous work in murine bone-marrow-derived macrophages (24, 25) and a recent study on Seahorse analysis of human monocyte-derived macrophage activation (29). A limitation of CENCAT with HPG over the original protocol is the requirement for methionine depletion before treatment, which requires specialized culture media and extra washing steps. Additionally, we observed very low levels of HPG incorporation in T cells and NK cells, indicating that these cells are metabolically quiescent under basal conditions, which is in accordance with previous research (30, 31), yet making it more difficult to accurately determine their metabolic dependencies. This was particularly evident for oligomycin treatment, which in some settings increased HPG incorporation compared to DMSO control, possibly due to metabolic adaptation through increased glycolytic flux (32). This phenomenon was also observed by Vogel et al. using puromycin, which they termed MITA (mitochondrial inhibition translation activation) (33). The occurrence of MITA could potentially lead to an underestimation of mitochondrial dependence.

As an alternative approach, we tested the ncAA βES, a novel threonine analog that is efficiently incorporated under native culture conditions (16). Our results showed that βES signal was higher than HPG in both naïve and activated T cells without depleting competing amino acids. Furthermore, we did not observe MITA in any tested cell types using βES. Application of CENCAT using βES in PBMCs replicated previously published metabolic signatures of immune cells. Upon activation, classical monocytes reportedly switch from a mitochondrial-dependent to a glycolytic profile (34), which could be reproduced by CENCAT. Furthermore, CD16^+^/CD14^-^ non-classical monocytes displayed higher levels of mitochondrial dependence compared to classical CD16^-^/CD14^+^ monocytes under basal conditions, in accordance with previous studies reporting increased mitochondrial activity and transcription of genes involved in mitochondrial respiration in non-classical monocytes (35, 36). Finally, activation of T cells increased their glycolytic capacity and/or glucose dependence as expected (37), particularly in CD45RA^+^ naïve and effector subsets.

Although studying circulating immune cell populations can yield valuable insights, it is important to acknowledge that they can differ significantly from their tissue-resident counterparts, which possess unique functions and metabolic profiles driven by microenvironmental imprinting. Until now, there has been a lack of suitable technologies to measure the metabolism of tissue-resident immune cells due to low cell availability and high sample complexity. By employing CENCAT on murine tissues, we could separate samples according to their tissue residency. Our results showed that the degree of βES incorporation was variable between tissues, with high levels detected in spleen and peritoneal immune cells, while βES signal was generally low in kidney samples. This is potentially related to the purity of the sample, as cells from the peritoneum and spleen are almost solely of leukocytic origin, whereas kidney samples still contain other cell types and biological material, which could interfere with βES uptake, even after CD45^+^ magnetic cell sorting. Metabolic dependencies of cDC2s and macrophage subsets were the main drivers of the separation between tissues. We further observed that tissue-resident cDC1 cells were generally more mitochondrially dependent than their cDC2 counterparts. Consistent with this finding, in vitro-generated cDC1s were reported to have an increased mitochondrial content and membrane potential compared to cDC2s (38). Among all tissues, neutrophils were found to be the most glycolytic of all studied cell types, which is congruent with current literature (39, 40). Within macrophage populations, kidney macrophages incorporated very low levels of βES, in line with their reported quiescent state during homeostasis (41). Furthermore, our results corroborate previous reports that peritoneal macrophages are highly mitochondrially dependent (42, 43). However, some of our results contradict a ranking of murine tissue macrophages based on their relative expression of OXPHOS-related genes in single-cell RNA sequencing datasets by Wculek and colleagues (44). For example, they showed alveolar macrophages among the highest-ranking subsets, while we found these cells to be relatively glycolytic among macrophage subsets. This indicates that OXPHOS gene signatures may not directly translate to functional preference of mitochondrial over glycolytic metabolism.

In summary, CENCAT is a promising technique for performing in-depth metabolic profiling of samples of varying complexity, spanning isolated cell types to ex vivo murine tissues. Our adaptation retains the benefits of the original protocol, including its independence of specialized equipment except for a flow cytometer, low cell number requirement, and non-necessity for cell purification. Compared to the original protocol, the use of ncAAs as a proxy for metabolic activity is preferable over puromycin due to their lack of toxicity. Based on our outcomes, βES is the superior ncAA for profiling complex samples containing cell types with variable metabolic activity. While tested here with a focus on immune cells, CENCAT is not limited by cell type or tissue and can also be applied in other research areas beyond immunometabolism.

## Declaration of interests

The authors declare no competing interests.

## Acknowledgements

HvdZ and RS are supported by the ZonMW grant TIMED (‘The right timing to prevent type 2 Diabetes’), project number 459001021.

---

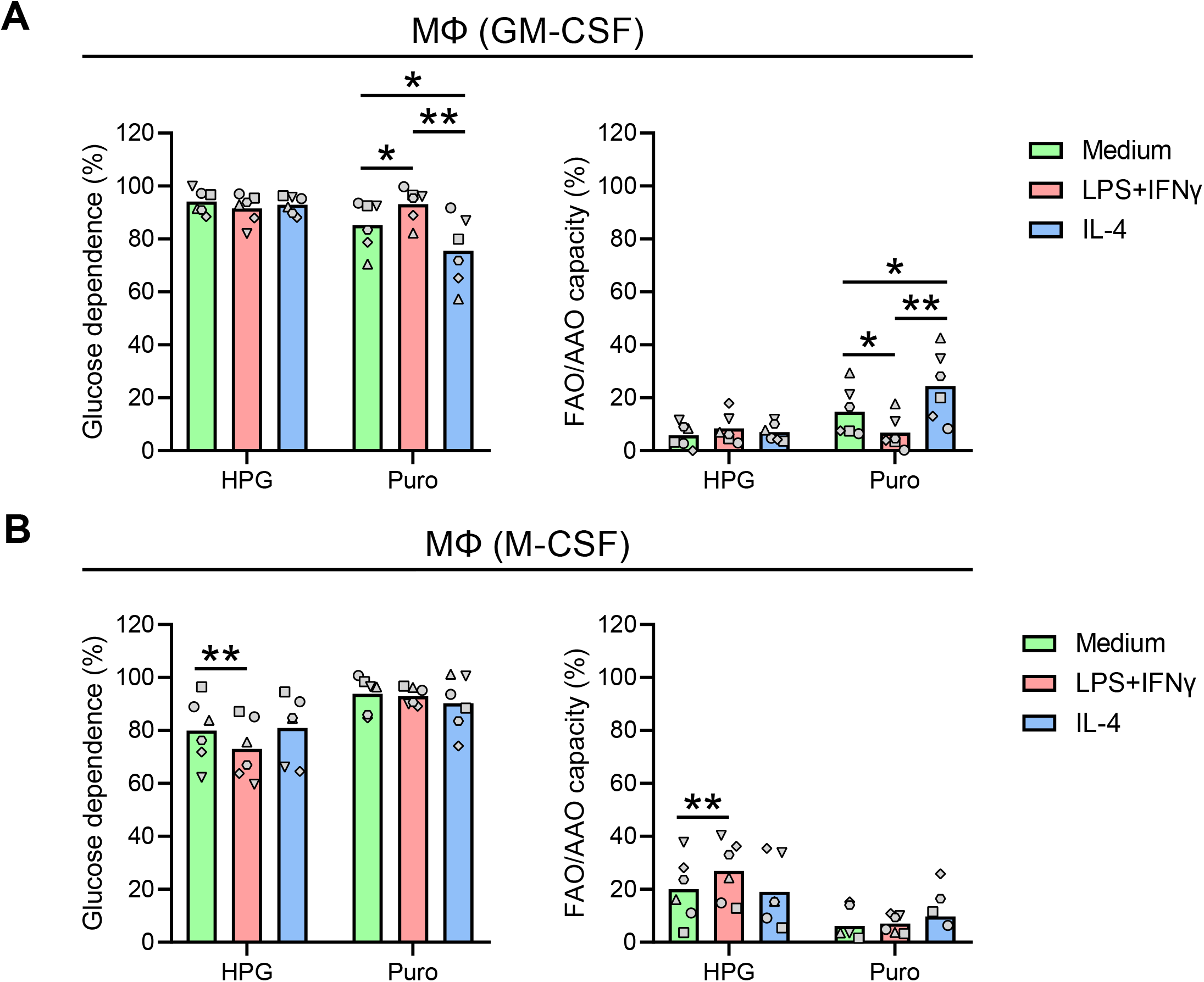

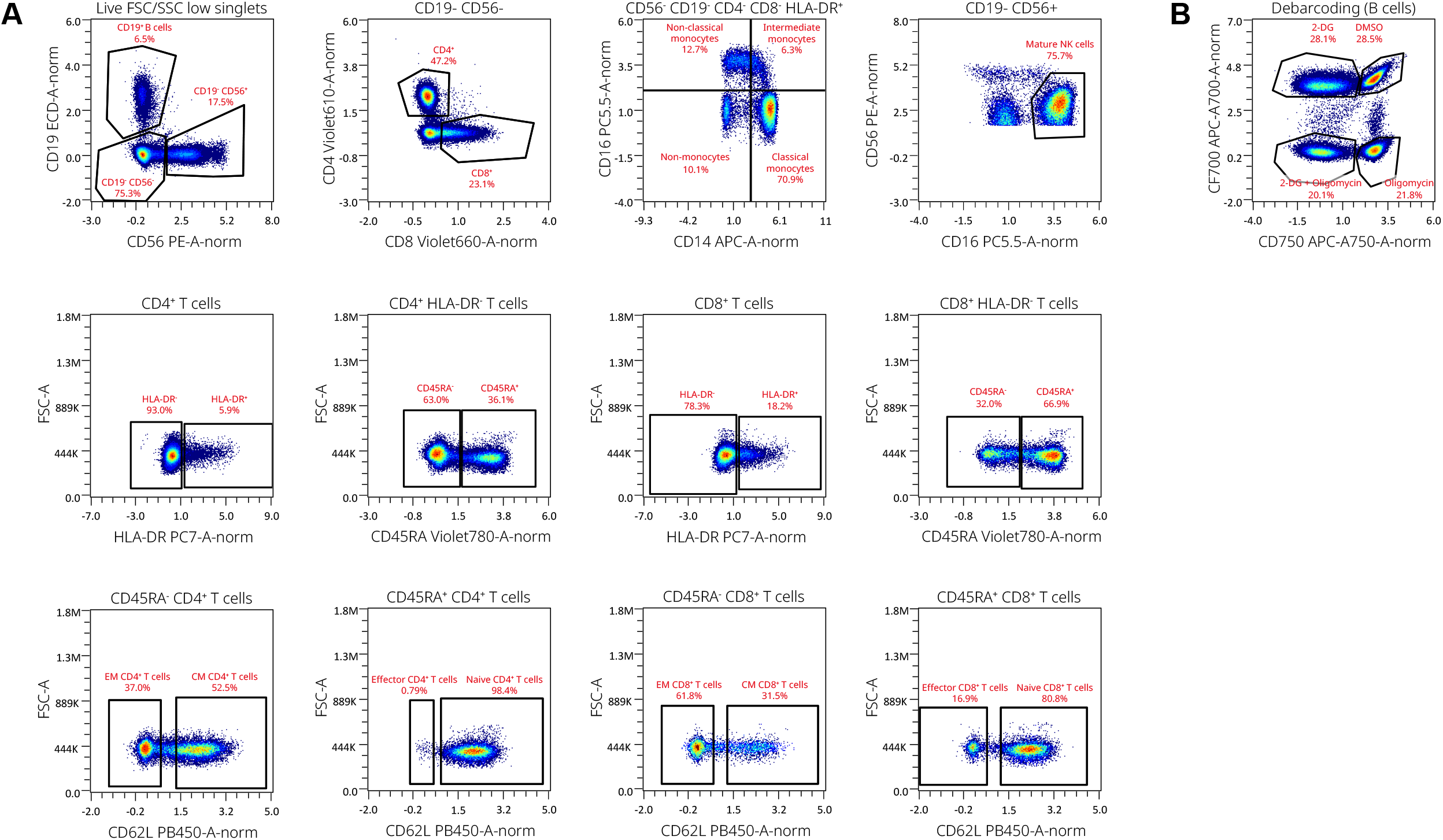

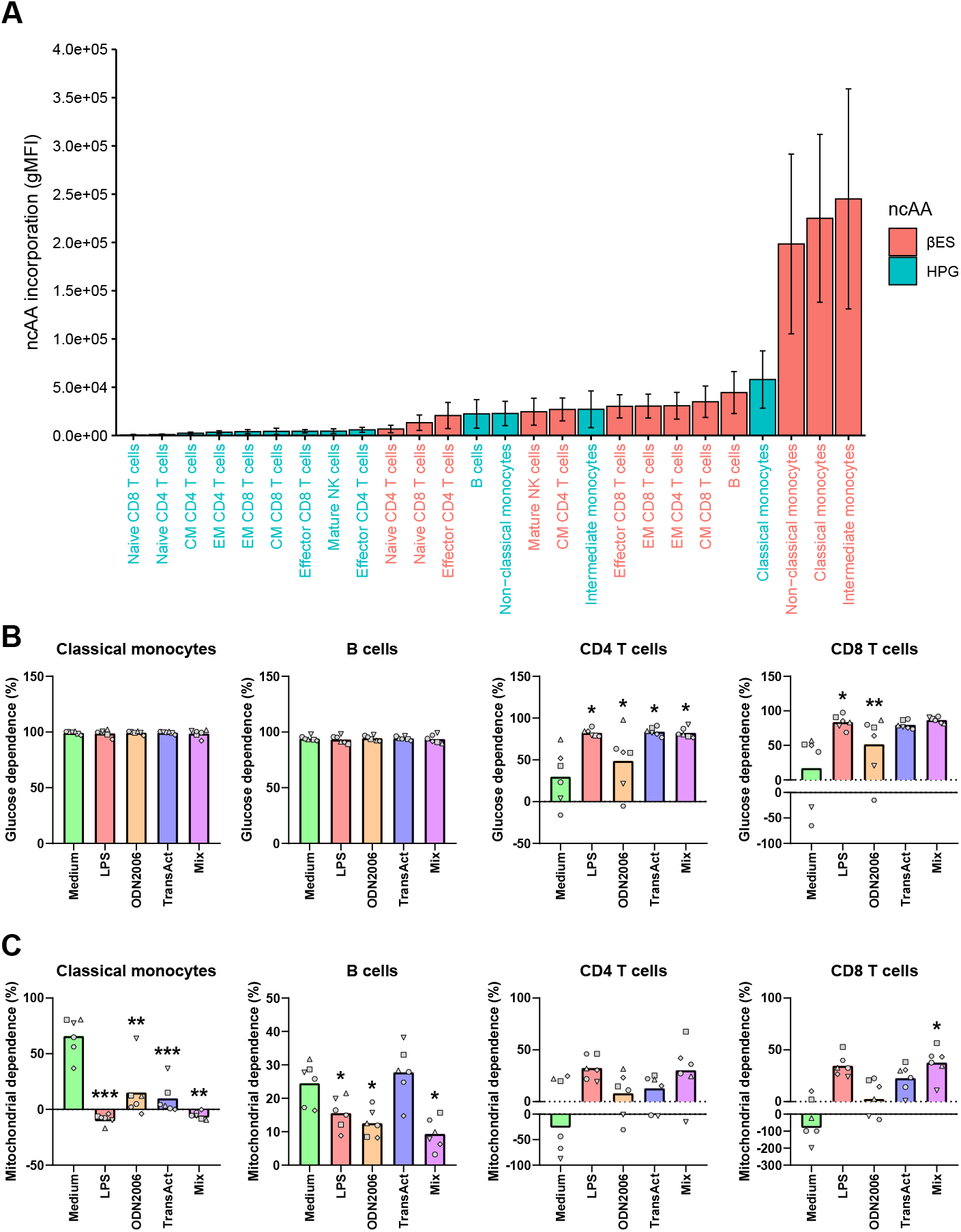

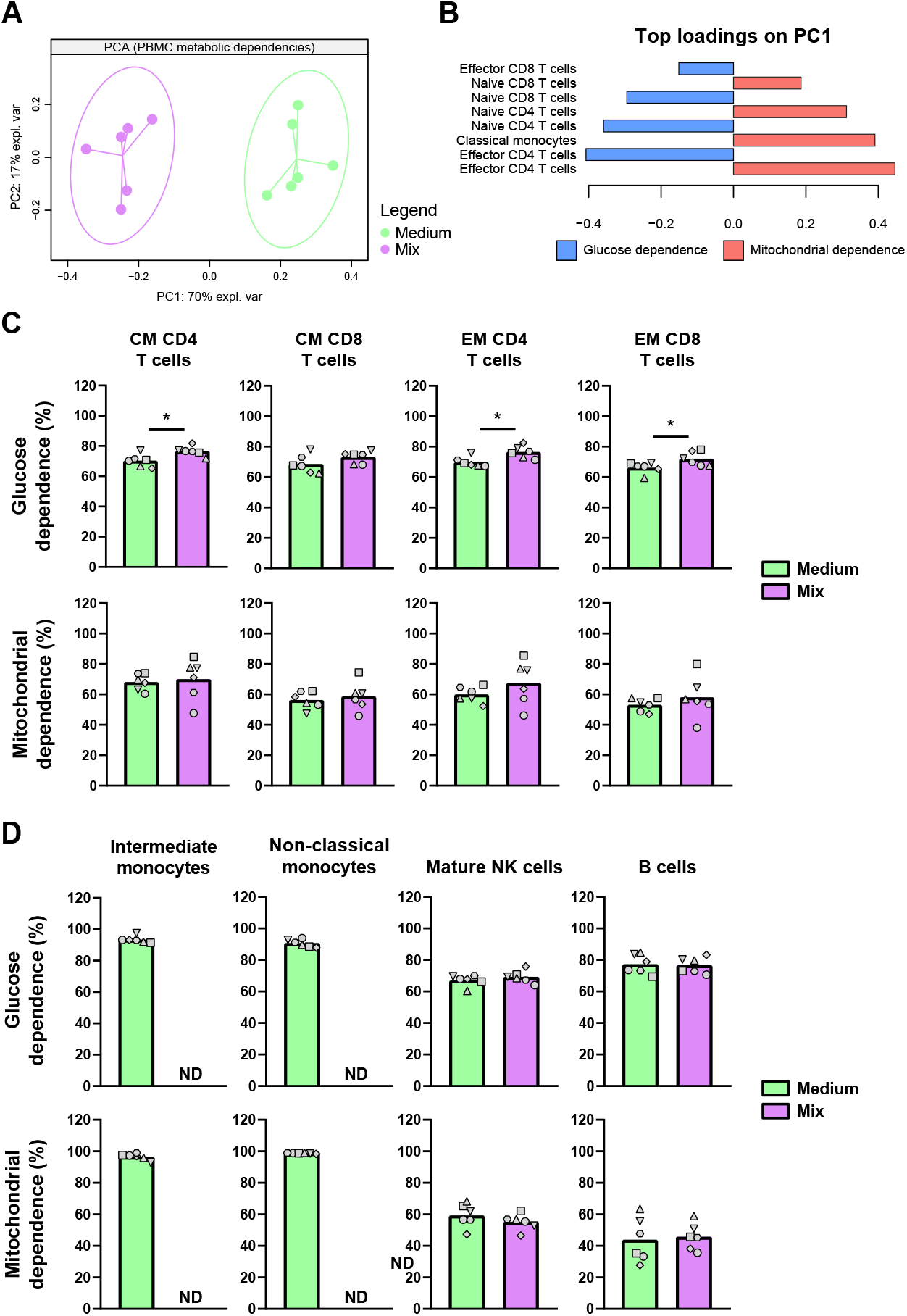

**Figure S5.**
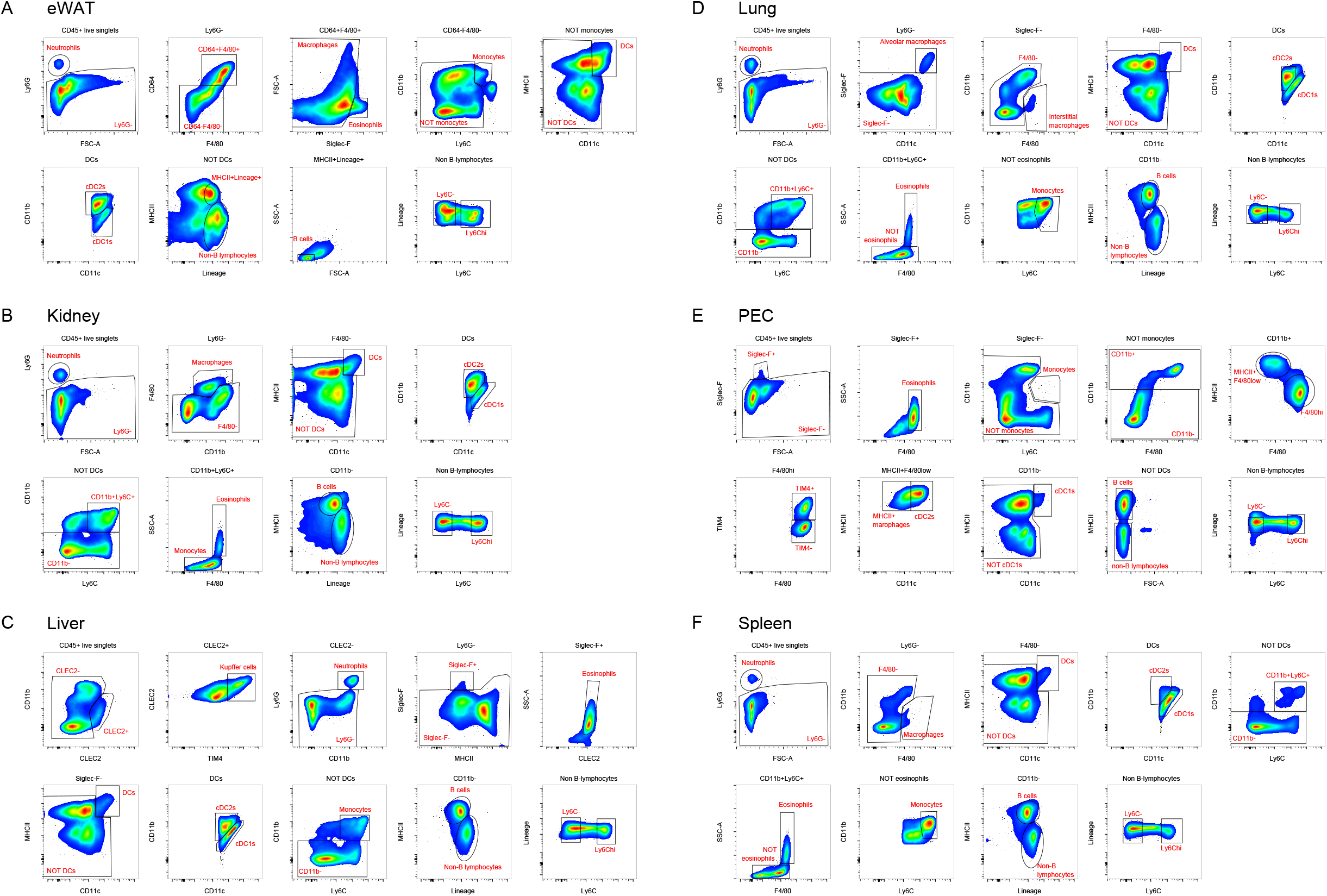

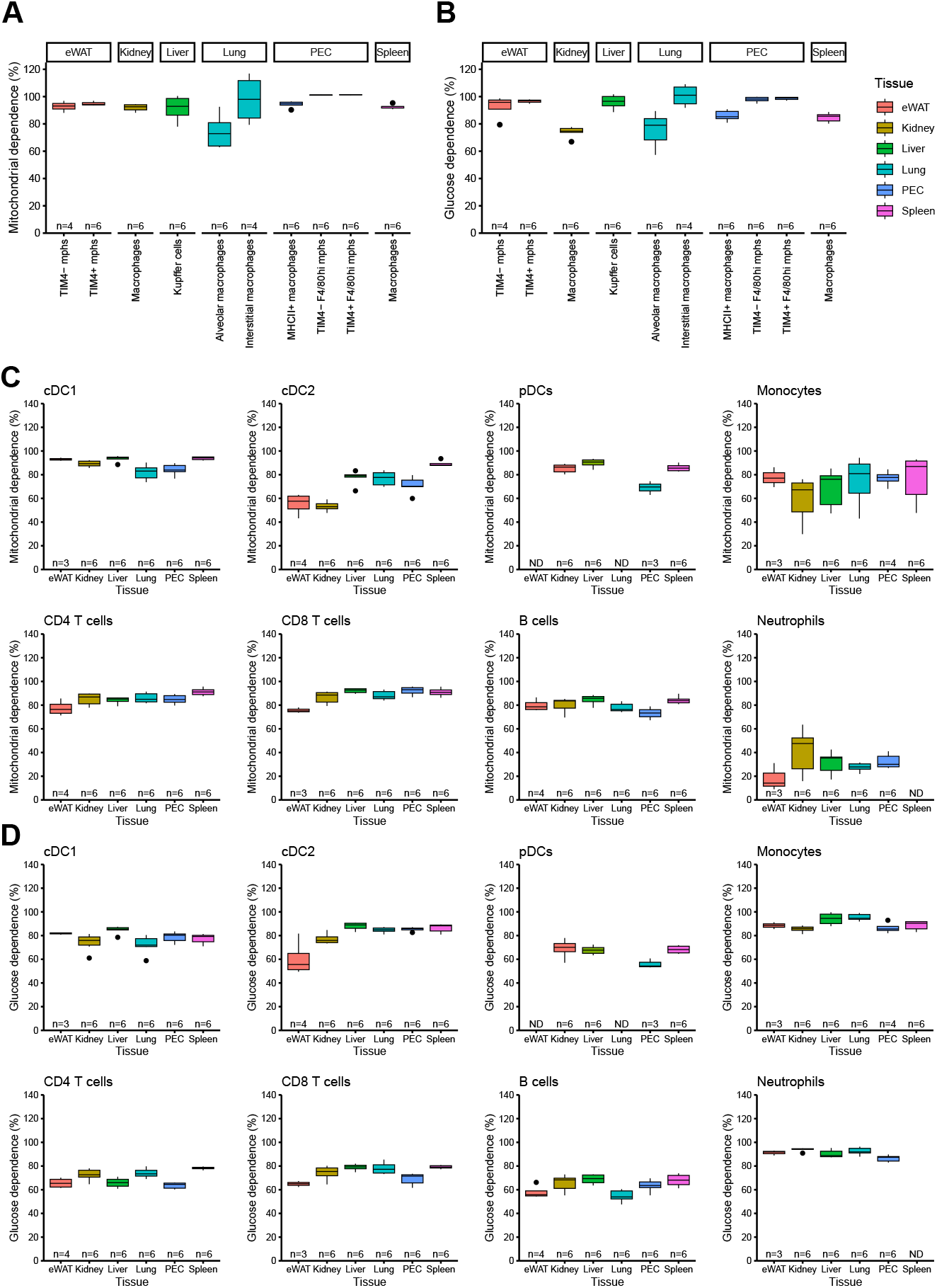

## Notes

### Competing Interest Statement

The authors have declared no competing interest.

### Summary of Updates

New data has been added to the manuscript regarding a novel non-canonical amino acid (bES). Based on these new insights, most main body and supplemental figures were updated/revised and the manuscript text was updated.

## References

1. Warburg O, Wind F, Negelein E. THE METABOLISM OF TUMORS IN THE BODY. J Gen Physiol. 1927;8(6):519–30.

2. Boutens L, Hooiveld GJ, Dhingra S, Cramer RA, Netea MG, Stienstra R. Unique metabolic activation of adipose tissue macrophages in obesity promotes inflammatory responses. Diabetologia. 2018;61(4):942–53.

3. Shirai T, Nazarewicz RR, Wallis BB, Yanes RE, Watanabe R, Hilhorst M, et al. The glycolytic enzyme PKM2 bridges metabolic and inflammatory dysfunction in coronary artery disease. The Journal of experimental medicine. 2016;213(3):337–54.

4. Kaymak I, Williams KS, Cantor JR, Jones RG. Immunometabolic Interplay in the Tumor Microenvironment. Cancer cell. 2021;39(1):28–37.

5. Verberk SGS, de Goede KE, Gorki FS, van Dierendonck X, Argüello RJ, Van den Bossche J. An integrated toolbox to profile macrophage immunometabolism. Cell reports methods. 2022;2(4):100192.

6. Argüello RJ, Combes AJ, Char R, Gigan JP, Baaziz AI, Bousiquot E, et al. SCENITH: A Flow Cytometry-Based Method to Functionally Profile Energy Metabolism with Single-Cell Resolution. Cell metabolism. 2020;32(6):1063-75.e7.

7. Oguma T, Ono T, Kajiwara T, Sato M, Miyahira Y, Arino H, et al. CD4(+)CD8(+) thymocytes are induced to cell death by a small dose of puromycin via ER stress. Cellular immunology. 2009;260(1):21–7.

8. Aviner R. The science of puromycin: From studies of ribosome function to applications in biotechnology. Computational and structural biotechnology journal. 2020;18:1074–83.

9. Marciano R, Leprivier G, Rotblat B. Puromycin labeling does not allow protein synthesis to be measured in energy-starved cells. Cell death & disease. 2018;9(2):39.

10. Dieterich DC, Link AJ, Graumann J, Tirrell DA, Schuman EM. Selective identification of newly synthesized proteins in mammalian cells using bioorthogonal noncanonical amino acid tagging (BONCAT). Proceedings of the National Academy of Sciences of the United States of America. 2006;103(25):9482–7.

11. Beatty KE, Xie F, Wang Q, Tirrell DA. Selective dye-labeling of newly synthesized proteins in bacterial cells. Journal of the American Chemical Society. 2005;127(41):14150–1.

12. Landgraf P, Antileo ER, Schuman EM, Dieterich DC. BONCAT: metabolic labeling, click chemistry, and affinity purification of newly synthesized proteomes. Methods in molecular biology (Clifton, NJ). 2015;1266:199–215.

13. Shen Y, Liu W, Zuo J, Han J, Zhang ZC. Protocol for visualizing newly synthesized proteins in primary mouse hepatocytes. STAR protocols. 2021;2(3):100616.

14. Pelgrom L, Davis G, O’Shoughnessy S, Kasteren SV, Finlay D, Sinclair L. QUAS-R: Glutamine (Q) Uptake Assay with Single cell Resolution reveals metabolic heterogeneity with immune populations. bioRxiv. 2022:2022.09.29.510040.

15. Su Hui Teo C, Serwa RA, O’Hare P. Spatial and Temporal Resolution of Global Protein Synthesis during HSV Infection Using Bioorthogonal Precursors and Click Chemistry. PLoS pathogens. 2016;12(10):e1005927.

16. Ignacio BJ, Dijkstra J, Mora N, Slot EFJ, van Weijsten MJ, Storkebaum E, et al. THRONCAT: metabolic labeling of newly synthesized proteins using a bioorthogonal threonine analog. Nature Communications. 2023;14(1):3367.

17. vanderZande HJP, Brombacher EC, Lambooij JM, Pelgrom LR, Zawistowska-Deniziak A, Patente TA, et al. Dendritic cell–intrinsic LKB1-AMPK/SIK signaling controls metabolic homeostasis by limiting the hepatic Th17 response during obesity. JCI Insight. 2023;8(11).

18. Liu Z, Gu Y, Shin A, Zhang S, Ginhoux F. Analysis of Myeloid Cells in Mouse Tissues with Flow Cytometry. STAR protocols. 2020;1(1):100029.

19. Le Cao K-A, Rohart F, Gonzalez I, Dejean S. mixOmics: Omics Data Integration Project. 2021.

20. Wickham H, Chang W, Henry L, Pedersen TL, Takahashi K, Wilke C, et al. ggplot2: Create Elegant Data Visualisations Using the Grammar of Graphics. 2023.

21. Wilke CO. cowplot: Streamlined Plot Theme and Plot Annotations for ggplot2. 2020.

22. van den Brand T. ggh4x: Hacks for ggplot2. 2023.

23. Van den Bossche J, O’Neill LA, Menon D. Macrophage Immunometabolism: Where Are We (Going)? Trends Immunol. 2017;38(6):395–406.

24. Krieg AM, Yi AK, Matson S, Waldschmidt TJ, Bishop GA, Teasdale R, et al. CpG motifs in bacterial DNA trigger direct B-cell activation. Nature. 1995;374(6522):546–9.

25. Hornung V, Rothenfusser S, Britsch S, Krug A, Jahrsdörfer B, Giese T, et al. Quantitative expression of toll-like receptor 1-10 mRNA in cellular subsets of human peripheral blood mononuclear cells and sensitivity to CpG oligodeoxynucleotides. Journal of immunology (Baltimore, Md : 1950). 2002;168(9):4531–7.

26. Kiick KL, Saxon E, Tirrell DA, Bertozzi CR. Incorporation of azides into recombinant proteins for chemoselective modification by the Staudinger ligation. Proceedings of the National Academy of Sciences of the United States of America. 2002;99(1):19–24.

27. Heieis GA, Patente TA, Almeida L, Vrieling F, Tak T, Perona-Wright G, et al. Metabolic heterogeneity of tissue-resident macrophages in homeostasis and during helminth infection. Nat Commun. 2023;14(1):5627.

28. Lacsina JR, Marks OA, Liu X, Reid DW, Jagannathan S, Nicchitta CV. Premature translational termination products are rapidly degraded substrates for MHC class I presentation. PloS one. 2012;7(12):e51968.

29. Hickman E, Smyth T, Cobos-Uribe C, Immormino R, Rebuli ME, Moran T, et al. Expanded characterization of in vitro polarized M0, M1, and M2 human monocyte-derived macrophages: Bioenergetic and secreted mediator profiles. PloS one. 2023;18(3):e0279037.

30. Chapman NM, Boothby MR, Chi H. Metabolic coordination of T cell quiescence and activation. Nature reviews Immunology. 2020;20(1):55–70.

31. Keppel MP, Saucier N, Mah AY, Vogel TP, Cooper MA. Activation-specific metabolic requirements for NK Cell IFN-γ production. Journal of immunology (Baltimore, Md : 1950). 2015;194(4):1954–62.

32. Hao W, Chang CP, Tsao CC, Xu J. Oligomycin-induced bioenergetic adaptation in cancer cells with heterogeneous bioenergetic organization. The Journal of biological chemistry. 2010;285(17):12647–54.

33. Vogel A, García González P, Argüello RJ. Measuring the Metabolic State of Tissue-Resident Macrophages via SCENITH. Methods in molecular biology (Clifton, NJ). 2024;2713:363–76.

34. Lachmandas E, Boutens L, Ratter JM, Hijmans A, Hooiveld GJ, Joosten LA, et al. Microbial stimulation of different Toll-like receptor signalling pathways induces diverse metabolic programmes in human monocytes. Nature microbiology. 2016;2:16246.

35. Zhao C, Tan YC, Wong WC, Sem X, Zhang H, Han H, et al. The CD14(+/low)CD16(+) monocyte subset is more susceptible to spontaneous and oxidant-induced apoptosis than the CD14(+)CD16(-) subset. Cell death & disease. 2010;1(11):e95.

36. Schmidl C, Renner K, Peter K, Eder R, Lassmann T, Balwierz PJ, et al. Transcription and enhancer profiling in human monocyte subsets. Blood. 2014;123(17):e90–9.

37. Frauwirth KA, Riley JL, Harris MH, Parry RV, Rathmell JC, Plas DR, et al. The CD28 signaling pathway regulates glucose metabolism. Immunity. 2002;16(6):769–77.

38. Kratchmarov R, Viragova S, Kim MJ, Rothman NJ, Liu K, Reizis B, Reiner SL. Metabolic control of cell fate bifurcations in a hematopoietic progenitor population. Immunology and cell biology. 2018;96(8):863–71.

39. Toller-Kawahisa JE, O’Neill LAJ. How neutrophil metabolism affects bacterial killing. Open Biol. 2022;12(11):220248.

40. Gainullina A, Mogilenko DA, Huang LH, Todorov H, Narang V, Kim KW, et al. Network analysis of large-scale ImmGen and Tabula Muris datasets highlights metabolic diversity of tissue mononuclear phagocytes. Cell reports. 2023;42(2):112046.

41. Wculek SK, Dunphy G, Heras-Murillo I, Mastrangelo A, Sancho D. Metabolism of tissue macrophages in homeostasis and pathology. Cellular & molecular immunology. 2022;19(3):384–408.

42. Woods PS, Kimmig LM, Meliton AY, Sun KA, Tian Y, O’Leary EM, et al. Tissue-Resident Alveolar Macrophages Do Not Rely on Glycolysis for LPS-induced Inflammation. American journal of respiratory cell and molecular biology. 2020;62(2):243–55.

43. Davies LC, Rice CM, Palmieri EM, Taylor PR, Kuhns DB, McVicar DW. Peritoneal tissue-resident macrophages are metabolically poised to engage microbes using tissue-niche fuels. Nat Commun. 2017;8(1):2074.

44. Wculek SK, Heras-Murillo I, Mastrangelo A, Mañanes D, Galán M, Miguel V, et al. Oxidative phosphorylation selectively orchestrates tissue macrophage homeostasis. Immunity. 2023;56(3):516-30.e9.

