## Supplemental Information for "In-depth immunometabolic profiling by measuring cellular protein translation inhibition via bioorthogonal noncanonical amino acid tagging (CENCAT)"

**Supplementary figure legends:**

**Figure S1: Glucose dependence and FAO/AAO capacity of GM-CSF & M-CSF macrophages.** Primary human macrophages (GM-CSF and M-CSF) were stimulated for 24 hours with culture medium (green), LPS + IFNγ (red), or IL-4 (blue) before SCENITH analysis using either HPG or puromycin (Puro) as substrates. (A) Glucose dependence (%) and FAO/AAO capacity (%) of GM-CSF macrophages. (B) Glucose dependence (%) and FAO/AAO capacity (%) of M-CSF macrophages. Data are displayed as mean percentages ± SD (n=6). Significance was tested by Two-Way ANOVA with Sidak correction for multiple testing. Individual donors are displayed by different symbols. * = *p* < 0.05, ** = *p* < 0.01, *** = *p* < 0.001

**Figure S2: Human PBMC 9-marker panel gating strategy.** (A) Representative gating of monocyte subtypes, B cells, NK cells, CD4 T cells and subtypes, and CD8 T cells and subtypes. (B) Representative debarcoding gating (B cells).

**Figure S3: CENCAT analysis of PBMCs using HPG.** PBMCs isolated from healthy blood donors (n=6) were stimulated for 2 hours with Medium control (green), LPS (red), ODN2006 (orange), TransAct (blue), or all stimuli combined (Mix, purple). (A) Relative incorporation (gMFI) of βES (red) and HPG (blue) in PBMC cell types under basal conditions (Medium control). (B) Glucose dependence (%) and (C) mitochondrial dependence (%) of classical monocytes, B cells, CD4 T cells and CD 8 T cells as determined by CENCAT using HPG as ncAA substrate. Significance was tested by Repeated One-Way ANOVA with Dunnett's multiple comparisons test. Bars indicate mean percentages and individual donors are displayed by different symbols. * = *p* < 0.05, ** = *p* < 0.01

**Figure S4: CENCAT analysis of PBMCs using βES.** PBMCs isolated from healthy blood donors (n=6) were stimulated for 2 hours with Medium control (green) or complete stimulation Mix (LPS+ODN2006+TransAct, purple). CENCAT was performed using βES as ncAA substrate. (A) PCA score plot based on metabolic dependencies of PBMCs. (B) Top loadings on PC1 of the PCA score plot. Measures of glucose dependence are represented by blue bars and mitochondrial dependence by red bars. Glucose dependence and mitochondrial dependence (%) of (C) CM CD4 T cells, CM CD8 T cells, EM CD4 T cells and EM CD8 T cells, (D) intermediate monocytes, non-classical monocytes, Mature NK cells and B cells. Significance was tested by paired t-test. Bars indicate mean percentages and individual donors are displayed by different symbols. ND = not detected. * = *p* < 0.05.

**Figure S5: Gating strategy mice tissues.** Representative gating of tissue-resident immune cell populations from (A) eWAT, (B) kidney, (C) liver, (D) lung, (E) PEC, and (F) spleen.

**Figure S6: Metabolic characteristics of murine tissue-resident immune cell populations.** The following tissues were isolated from male C57BL/6J mice and subjected to CENCAT analysis: eWAT (red), kidney (yellow), liver (green), lung (cyan), PEC (blue) and spleen (pink). (A) Boxplots of glucose dependence (%) and (B) mitochondrial dependence (%) of tissue-resident macrophage populations. (C) Mitochondrial dependence (%) of cDC1s, cDC2s, pDCs, monocytes, CD4 T cells, CD8 T cells, B cells, and neutrophils from all six tissues. (D) Mitochondrial dependence (%) of cDC1s, cDC2s, pDCs, monocytes, CD4 T cells, CD8 T cells, B cells, and neutrophils from all six tissues. Amount of samples (n) is indicated for each boxplot. ND = not detected.
